# Dissecting the Metabolic Reprogramming of Maize Root under Nitrogen Limiting Stress Condition

**DOI:** 10.1101/2021.04.30.442195

**Authors:** Niaz Bahar Chowdhury, Wheaton L. Schroeder, Debolina Sarkar, Nardjis Amiour, Isabelle Quilleré, Bertrand Hirel, Costas D. Maranas, Rajib Saha

## Abstract

The growth and development of maize (Zea mays L.) largely depends on its nutrient uptake through root. Hence, studying its growth, response, and associated metabolic reprogramming to stress conditions is becoming an important research direction. A genome-scale metabolic model (GSM) for the maize root was developed to study its metabolic reprogramming under nitrogen-stress condition. The model was reconstructed based on the available information from KEGG, UniProt, and MaizeCyc. Transcriptomics data derived from the roots of hydroponically grown maize plants was used to incorporate regulatory constraints in the model and simulate nitrogen-non-limiting (N−) and nitrogen-deficient (N−) conditions. Model-predicted result achieved 70% accuracy comparing to the experimental direction change of metabolite levels. In addition to predicting important metabolic reprogramming in central carbon, fatty acid, amino acid, and other secondary metabolism, maize root GSM predicted several metabolites (e.g., L-methionine, L-asparagine, L-lysine, cholesterol, and L-pipecolate) playing critical regulatory role in the root biomass growth. Furthermore, this study revealed eight phosphatidyl-choline and phosphatidyl-glycerol metabolites which even though not coupled with biomass production played a key role in the increased biomass production under N-. Overall, the omics-integrated-GSM provides a promising tool to facilitate stress-condition analysis for maize root and ultimately engineer better stress-tolerant maize genotypes.

**Summary:**

- The growth and development of maize *(Zea mays* L.) largely depends on its nutrient uptake through root. Hence, studying its growth, response, and associated metabolic reprogramming to stress conditions is becoming an important research direction.
- A genome-scale metabolic model (GSM) for the maize root was developed to study its metabolic reprogramming under nitrogen-stress condition. The model was reconstructed based on the available information from KEGG, UniProt, and MaizeCyc.
- Transcriptomics data derived from the roots of hydroponically grown maize plants was used to incorporate regulatory constraints in the model and simulate nitrogen-non-limiting (N^+^) and nitrogen-deficient (N^−^) conditions. Model-predicted result achieved 70% accuracy comparing to the experimental direction change of metabolite levels. In addition to predicting important metabolic reprogramming in central carbon, fatty acid, amino acid, and other secondary metabolism, maize root GSM predicted several metabolites (e.g., L-methionine, L-asparagine, L-lysine, cholesterol, and L-pipecolate) playing critical regulatory role in the root biomass growth. Furthermore, this study revealed eight phosphatidyl-choline and phosphatidyl-glycerol metabolites which even though not coupled with biomass production played a key role in the increased biomass production under N^−^.
- Overall, the omics-integrated-GSM provides a promising tool to facilitate stress-condition analysis for maize root and ultimately engineer better stress-tolerant maize genotypes.

## Introduction

Maize (*Zea mays* L.) is considered as one of the major sources of food for a large portion of world population (Rouf Shah et al., 2016). According to the International Grains Council, global maize consumption will climb to new peaks in the coming years, as well as the usage of maize as food is forecasted to expand (International Grains Council, 2019). Nutrients that are uptaken by roots are essential for maize plant growth and limited nutrient supply can stymie plant growth along with discernable phenotypical changes (Hu & Chu, 2020). Among these nutrients, nitrogen (N) plays a key role in plant growth. N limitation frequently reduces crop growth and yield, and contributes to a variety of phenotypical changes including expanded root architecture (Mager & Ludewig, 2018), increased root biomass (Mardanov et al., 1998), and root exudate profiles (Frey et al., 2009; Neal et al., 2012). Although many experimental studies are available that study the metabolism associated with nitrogen starvation of maize root (Krapp et al., 2011; Mager & Ludewig, 2018; Schlüter et al., 2012; Tschoep et al., 2009), they are primarily focused on probing specific class of metabolites, such as amino acid or fatty acids. But capturing the aggregate effect of the maize root physiology on biomass production under N^−^ is essential to understand plant-wide reprogramming of its metabolism. This sort of reprogramming has been reported as the hallmark of metabolism which allows any system to adapt to the changes to external conditions (Medina, 2020). Thus, in order to capture this metabolic reprogramming of maize root under N^−^ in an integrated manner, developing system biology approaches will allow to have a dynamic picture of plant adaptation to nitrogen deficiency.

To develop such system biology approach, genome-scale metabolic models (GSMs) have been widely used either at the organ or the whole plant level (Shaw & Cheung, 2020). A GSM captures most of the known metabolic reactions within a biological system (i.e., a prokaryotic/eukaryotic organism or an organ/a group of organs of a higher-order organism) and can predict the flux of reactions by implementing techniques such as flux balance analysis (FBA) (Orth et al., 2010), flux variability analysis (FVA) (Mahadevan & Schilling, 2003), and parsimonious FBA (pFBA) (Lewis et al., 2010). Typically, reaction flux is predicted by solving an optimization problem through maximizing biomass production. In higher plants, the first GSM was reconstructed for Arabidopsis more than a decade ago (Poolman et al., 2009), thus setting up a new direction of research with the aid of systems biology. Once the complete genome sequences of more plants became available, metabolic reconstruction picked up pace, giving a range of published plant GSMs, including crops such as maize (*Zea mays* L.) (Saha et al., 2011), rice (*Oryza sativa L.*) (Chatterjee et al., 2017; Poolman et al., 2013), rapeseed (*Brassica napus L.*) (Pilalis et al., 2011), and model plants such as Arabidopsis (*Arabidopsis thaliana*) (Cheung et al., 2013; Dal’Molin et al., 2010; Poolman et al., 2009), and *Setaria viridis L.* (Shaw & Maurice Cheung, 2019). Although these GSMs provided reconstruction of the whole plant metabolism, organ-specific GSMs including those of rapeseed embryo (Hay & Schwender, 2011), barley seed (Grafahrend-Belau et al., 2009), Arabidopsis leaf (Arnold & Nikoloski, 2014), as well as for maize leaf (Seaver et al., 2015; Simons et al., 2014), endosperm (Seaver et al., 2015), and embryo (Seaver et al., 2015) were developed in parallel. These organ-specific GSMs provided better resolution of the metabolism in specialized tissues, which can be probed further for useful insights on whole plant physiology. Up to now, there is no maize root-specific GSM available to investigate its metabolism/associated reprogramming under stress conditions such as mineral nutrient deficiency and derive new biological insights that can experimentally be tested.

An important challenge to further sharpen model predictions and, thereby, decipher meaningful biological information from a GSM is to integrate environment/condition-specific ‘omics’ data into it. Although there is a paucity of environment specific ‘omics’ data (i.e., gene expression, protein abundance, and metabolite level) for maize root, their availability would help constrain the solution space and thus improving final model predictions. Proteomics and transcriptomics data can be used to apply flux constraints on corresponding reactions determined by gene-protein-reaction (GPR) associations via switch (e.g., GIMME (Becker & Palsson, 2008), iMAT (Zur et al., 2010), and MADE (Jensen & Papin, 2011)) or valve (e.g., E-Flux (Colijn et al., 2009) and PROM (Chandrasekaran & Price, 2010)) approach. While the switch approaches have a binary nature, resulting in all-or-nothing posture to constraining reactions which may not reflect the actual condition, valve approaches provide more flexibility in constraining the solution space by incorporating the gene expression/protein abundance data as reaction flux constraints. In addition to incorporating these reaction regulations, model-generated metabolite levels can be qualitatively compared with experimental metabolite measurements, thus further characterizing the state of metabolism. To this end, flux-sum is used as a proxy of the metabolite pool size (Chung & Lee, 2009). Although fast or slow kinetics may diminish or expand the size of the pool, without changing corresponding flux.

In this work, maize plants were grown in hydroponic culture system under N^+^ and N^−^ and used to generate root transcriptomics and metabolomics datasets. In parallel, a GSM was built for roots in order to incorporate transcriptomic and metabolic profiling data into the model to simulate the metabolic reprogramming of maize roots under N^−^. The model was first validated using a study from literature (Walton et al., 2016) on the effect of synthetic strigolactone, rac-GR24, on the root and its associated metabolic reprogramming. Next, the transcriptomics data was used to implement regulations in the GSM and subsequently to simulate N^+^ and N^−^. When compared with metabolites, the model-predicted results achieved 70% accuracy concerning the reprogramming of central carbon (C), fatty acid, amino acid, and secondary metabolism. In addition, the GSM allowed the prediction of regulatory roles of several metabolites in root biomass production, such as L-methionine, L-asparagine L-lysine, cholesterol, and L-pipecolate. Eight phosphatidyl-choline and phosphatidyl-glycerol metabolites not coupled to biomass production under N^+^, were found to play a seminal role in the increased biomass growth under N^−^. Going forward, we expect that this maize root GSM will provide a powerful tool to study the effect of other abiotic and biotic stresses such as phosphate deficiency, salinity, heat stress, draught, heavy metal stresses, rot disease, and interactions with beneficial microorganisms.

## Materials and Methods

### Plant material

Plant material (*Zea mays* L., line B73) for RNA extraction, biomass components and metabolite measurements were grown in a hydroponic culture system under controlled conditions. To avoid heterogeneity in the germination time, imbibition of the seeds was performed at 6°C in the dark for 3 days in petri dished containing filter paper moisten with sterile distilled water. Seedlings were then transferred onto sand and watered daily on a nutrient solution containing 5.6 mM K^+^, 3.4 mM Ca^2+^, 0.9 mM Mg^2+^, 0.9 mm H_2_PO_4_-, and 21.5 mM Fe (Sequestrene Ciba-Geigy, Basel, 23 mM B, 9 mM Mn, 0.30 mM Mo, 0.95 mM Cu, and 3.50 mM Zn. Nitrogen was supplied as 1 mM KNO_3_-. After 1 week when two to three leaves had emerged, plants were randomly placed on a 130-L aerated hydroponic culture unit containing a complete nutrient solution (N^+^) containing 5mM NO_3_-(Coïc & Lesaint, 1971). The complete nutrient solution (N^+^) contained 5 mM K^+^, 3 mM Ca^2+^, 0.4 mM Mg^2+^, 1.1 mM H2PO_4_-, 1 mM SO_4_2^−^, 1.1mM Cl^−^,21.5 μM Fe^2+^ (Sequestrene; Ciba-Geigy, Basel, Switzerland), 23 μM B^3+^, 9 μM Mn^2+^, 0.3 μM Mo^2+^, 0.95 μM Cu^2+^ and 3.5 μM Zn^2+^. For growing plants under N^−^, NO_3_-was supplied as 0.1mM KNO_3_-, a N concentration that has previously been shown to provide N-deficiency stress for most plant species (Amiour et al., 2012; Hirel et al., 2005; Tercé-Laforgue et al., 2004). The N^+^ and N^−^ nutrient solutions were replaced daily. The experiment was performed in triplicate for each N concentration in the nutrient solution in separate hydroponic units placed side by side. The six hydroponic culture units were kept 18 days in a controlled environment chamber in 2014 (May 18–June 2014). Three plants in each hydroponic unit were pooled making three replicates for each of the two N feeding conditions. Plants were harvested at the 6 to 7 leaf stage between 9 to 12 am and separated into shoots and roots. The samples were immediately placed in liquid N_2_ and then stored at −80°C until further analysis.

### Gene expression profiles using maize cDNA microarrays

RNA preparation, transcript expression profiles and statistical analysis of the data were performed essentially as described in literature (Amiour et al., 2012).

### Metabolite extraction and analyses

Frozen root material previously stored at −80°C was used for metabolite extraction. C. 100 mg of the powder was extracted in 1 ml of 80% ethanol/20% distilled water for an hour at 4°C. During extraction, the samples were continuously agitated and then centrifuged for 5 min at 15,000 rpm. The supernatant was removed, and the pellet was subjected to a further extraction in 60% ethanol and finally in water at 4°C, as described above. All supernatants were combined to form the aqueous alcoholic extract. Total N content of 25 mg of frozen root material was determined in an elemental analyzer using the combustion method of Dumas (Flash 2000, Thermo Scientific, Cergy-Pontoise, France). RNA, DNA, amino acids, proteins, lipids, starch, soluble carbohydrates, and cell wall carbohydrates content were quantified according to the different protocols from literature (Simons et al., 2014).

### Metabolome analysis

All steps were adapted from the original protocol (Fiehn, 2006) following the procedure described in literature (Amiour et al., 2012). Detail experimental procedure can be found in Supplementary Information Notes S1.

### Model reconstruction

The primary set of reactions *J*_Primary_ (from a combination of gene, protein, and reaction information from available public databases such as Kyoto Encyclopedia of Genes (Kanehisa et al., 2014), UniProt (Bateman, 2019), and MaizeCyc (Monaco et al., 2013)), vascular tissue transporters *Vt*, and nutrients *J*_Nutrients_ supplied to maize root were combined to form the draft model for maize root. Genes with an expression level above the cut-off defined by Downs et al. (i.e., 7.644) were categorized as highly expressed. Genes that were expressed at a level below the cut-off were categorized as lowly expressed. When building maize root model, the nutrients included those from the soil and those metabolites that can be imported from the vascular tissue. Inactive reactions, determined by using the gene expression data and gene-protein-reaction (GPR) associations, were placed into a set termed *J*_Inactive_. Finally, spontaneous reactions and reactions with a GPR relationship that only contain genes in the “always lowly expressed” set were placed into the set *J*_NotMeasured_. To ensure that maize root biomass reaction was not blocked given the available nutrients, a GapFill step (Satish Kumar et al., 2007) was performed for the maize root. The following algorithm was used to determine the minimum number of reactions added with a preference of adding reactions from the not measured set over reactions in the inactive set (Equation 1):

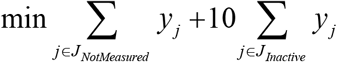

*Subjecte to*:

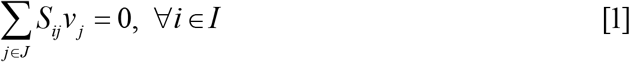

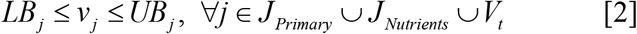

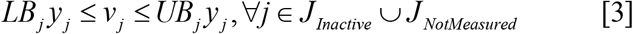

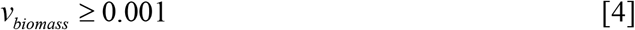

Here *S*_*ij*_ is the stoichiometric coefficient of metabolite *i* in reaction *j* and *v*_*j*_ is the flux through reaction *j.* Sets *I* and *J* include all metabolites and reactions known to occur within maize, respectively. The lower bound, *LB_j_*, and upper bound, *UB_j_*, of reaction *j* (Equations 2 and 3) are sufficiently small and large and are determined by each reaction’s directionality based on thermodynamic constraints. The binary variable, *y_j_*, is equal to 1 if the reaction is added to the model and 0 otherwise (Equation 3). Finally, a small amount of flux is forced through the biomass equations (Equation 4) to ensure biomass is not blocked. From this algorithm, a set of reactions, *J*_Secondary_, was added to the maize root models to ensure flux through the biomass reaction is possible.

A second GapFill step was performed to activate as many reactions in the primary set as possible for maize root. The GapFill algorithm displayed above was completed successively for each reaction that did not carry flux in the primary set by modifying the model constraints in the following manner. First, equation 2 was extended to encompass the secondary set of reactions to account for their inclusion in the maize root model. Second, equation 4 was applied to reactions of interest rather than the biomass reaction. Reactions from *J*_NotMeasured_ were added as needed, however reactions from *J*_Inactive_ were only included in the final model if their addition allowed ten previously blocked reactions from the active set to carry flux. This was done to ensure that reactions with genes that are lowly expressed in the maize root based on transcriptomic data were added only if they allow for flux through multiple reactions in the primary set.

### Eliminating thermodynamically infeasible cycles through flux variability analysis

To identify thermodynamically infeasible cycles in the model, FVA (Mahadevan & Schilling, 2003) was used by turning all the nutrient uptakes to the cell off. The formulation is as follows.

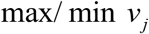

*Subject to*:

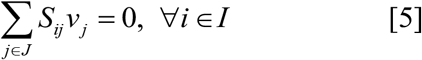

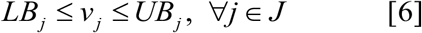

FVA maximizes and minimizes each of the reaction fluxes subject to mass balance and environmental, and any artificial (i.e., biomass threshold) constraints. The reaction fluxes which hit either the lower or upper bounds, were defined as unbounded reactions, and were grouped as a linear combination of the null basis of their stoichiometric matrix. These groups are indicative of possible thermodynamically infeasible cycles. To eliminate the cycles, duplicate reactions were removed, lumped reactions were turned off, or reactions were selectively turned on/off based on available cofactor specificity information.

### Incorporation of transcriptomics data with the model through E-Flux

E-Flux is an extension of Flux Balance Analysis (FBA) that infers a metabolic flux distribution from transcriptomic data (Brandes et al., 2012; Colijn et al., 2009). The rationale behind E-Flux is, given a limited translational efficiency and a limited accumulation of enzyme over the time, the mRNA level can be used as an approximate upper bound on the maximum number of metabolic enzymes, and corresponding reaction rates. The standard FBA involved solving the following linear optimization problem:

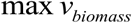

*Subject to*:

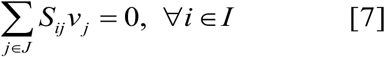

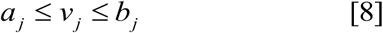

where vector *v* represents a particular flux configuration, *S* is the stoichiometric matrix, and *a*_*j*_ and *b*_*j*_ are the minimum and maximum allowed fluxes through reaction *j*. It was assumed that a set of expression measurements for some or all of the genes associated with the reactions in *S* were available. The core E-Flux method chooses the maximum flux, *b*_*j*_, for the *j*^*th*^ reaction according to a function of the expression of gene *j* and associated genes:

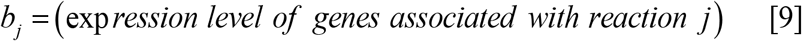

Expression level of each reaction was determined through its Gene-Protein-Reaction (GPR) relationship. If the reaction catalyzed by the corresponding enzyme was reversible then *a*_*j*_ = −*b*_*j*_, otherwise *a*_*j*_ = 0.

### Metabolite pool size calculations through flux-sum analysis

Metabolite turnover rates were determined based on the flux-sum analysis (FSA) method (Chung & Lee, 2009) and compared with the metabolomic data. The flux-sum is a measure of the amount of flow through the reactions associated with either the production or consumption of the metabolite. The range of the flux-sum or the flow through of each metabolite with experimental measurements was maximized/minimized as follows:

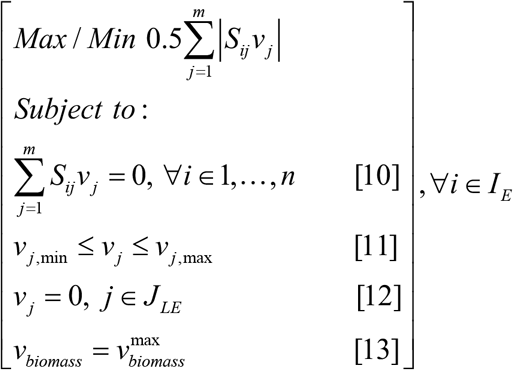

Here, set *I*_*E*_ represents the set of metabolites with experimental measurements and set *J*_*LE*_ represents reactions with statistically lower expression of gene transcripts and/or proteins. The formulation was run in an iterative manner for each metabolite with experimental measurements and repeated for each individual condition. By linearizing the objective function, the resulting formulation is a mixed-integer linear programming problem. Therefore, the basic idea was to determine the range of the flux-sum of a metabolite (for which metabolomic data are available) under a given condition by switching off reaction fluxes corresponding to gene transcripts and/or proteins with lower expression level (equation 12). The flux-sum ranges were determined at the maximum biomass for the condition as displayed in equation 13. Predictions were made only when the flux-sum ranges did not overlap between the background condition and the condition to be compared. In this way, the compartment-specific predictions of the flux-sum ranges were compared with tissue-specific experimental measurements. As GSM is a steady state model, hence diminishing or expanding pool size based on fast or slow reaction kinetics is not a possibility here.

### Use of BLASTp algorithm to find homologous genes

Arabidopsis genes sequence data was obtained from the Tair database (http://www.arabidopsis.org/). To identify Arabidopsis gene homologs in maize, systematic bidirectional BLASTp (https://blast.ncbi.nlm.nih.gov/Blast.cgi?PAGE=Proteins) searches were performed against the NCBI non-redundant database using the sequences of Arabidopsis genes. The screening criteria was E value 10^−10^. In case of multiple homologous genes for maize, genes were chosen based on the higher Max score.

### Simulation software

The General Algebraic Modeling System (GAMS) version 24.7.4 with IBM CPLEX solver was used to run FBA and FVA, E-Flux, and FSA algorithm on the model. Each of the algorithms was scripted in GAMS and then run on a Linux-based high-performance cluster computing system at the University of Nebraska-Lincoln.

## Results and Discussions

### Development and validation of maize root model

A GSM of maize root was reconstructed using a combination of gene, protein, and reaction information from available public databases such as Kyoto Encyclopedia of Genes (Kanehisa et al., 2014), UniProt (Bateman, 2019), and MaizeCyc (Monaco et al., 2013). The draft model was curated as described in the Materials and Methods section. Fig. 1 also depicts how model reconstruction and refinement carried out. The model also included the metabolites that are transported through the phloem and xylem tissues supported by literature evidence (Yesbergenova-Cuny et al., 2016). The phloem and xylem tissues were combined, for simplicity, in the model and are referred to as the vascular tissue. Specific pathways such as sphingolipid pathway, benzoxazinoid pathway, linoleic acid pathway, β-alanine pathway, and flavonoid biosynthesis were introduced based on genomic annotation and experimental evidence from literature as described in the sub-sequent sections. The reconstructed model contains 6389 genes, 4002 reactions, and 4461 metabolites distributed across six intracellular compartments, namely, cytosol, plastid, mitochondria, peroxisome, plasma membrane, vacuole, and inner mitochondrial matrix. Biomass accumulation is represented in the metabolic model by a root-specific biomass. Within the biomass reaction, the stoichiometric coefficients represent the proportion of each biomass component ensuring that the overall biomass molecular weight is 1 gm mol^-1^ for proper component balance (Chan et al., 2017). Growth and non-growth associated ATP maintenance (GAM and NGAM, respectively) levels were based on available measurements for maize root (Doncheva et al., 2006; Roberts et al., 1985).

**Fig. 1:**
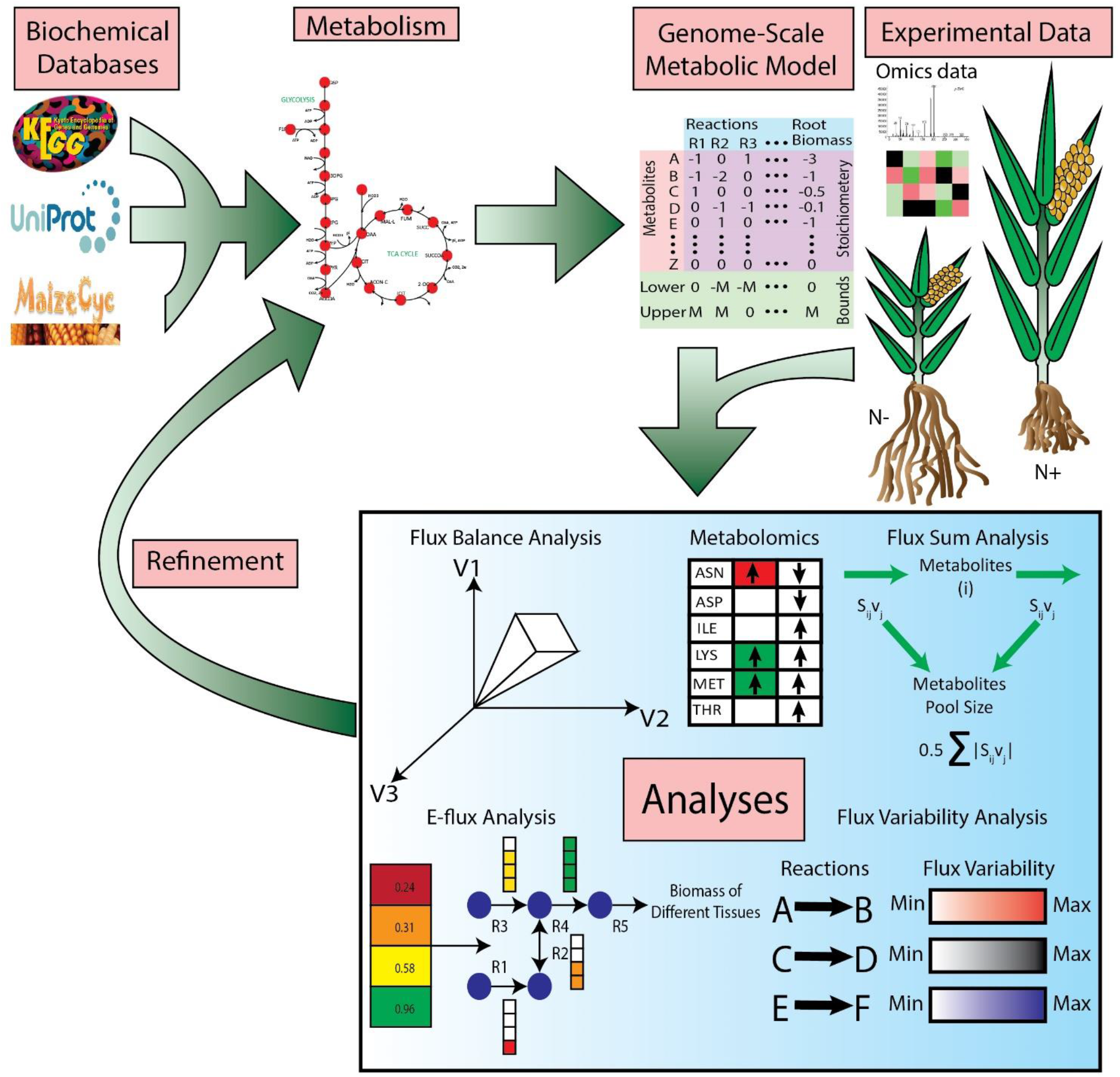
Model reconstruction, omics data integration, and model refinement. The maize root genome-scale metabolic model can be found in Supplementary Information Table S1. Experimental transcriptomics and metabolomics data can be found in Supplementary Information Table S2.

In order to validate the model and to ensure that it could simulate biologically relevant situations, a case study involving strigolactones was selected. Strigolactones, a group of carotenoid-derived terpenoid lactones, can regulate root architecture, act as phytohormones and rhizosphere signals (Guan et al., 2012; Walton et al., 2016). Strigolactones also promote the elongation of seminal/primary roots and adventitious roots and they repress lateral root formation (Sun et al., 2016). Given the importance of strigolactones, the key question was if the maize root model could recapitulate the manifestation of these effects in terms of metabolic changes. To our knowledge, no ‘omics’ data is available for maize regarding the effect of strigolactones. Thus, the relevant information was gathered from a study performed in *Arabidopsis thaliana* (Walton et al., 2016) in which the impact of the strigolactone analog *rac*-GR24 was elucidated on the root proteome of the wild type (WT) and the signaling mutant *more axillary growth 2 (max2*). The study revealed a clear MAX2-dependent *rac*-GR24 response, which indicated an increase in abundance of enzymes involved in the flavonol biosynthesis. This abundance of enzyme in flavonol biosynthesis was reduced in the *max2–1* mutant. Details of the experimental design and screening of these genes can be found in the original paper (Walton et al., 2016). To identify the homologous genes encoding the corresponding proteins in maize, a bidirectional BLASTp search was conducted. Later, the abundance of those proteins was used to incorporate reaction flux regulations via GPR in the maize root model for predicting the response of the maize root to the synthetic strigolactone *rac*-GR24.

As the number of homologous genes with measured protein levels were low in numbers, most of the metabolic pathways between WT and *rac*-GR24 stimulated maize root showed similar flux ranges. However, the maize root model was able to predict the reduced reaction flux in flavonoid biosynthesis pathway and phenylpropanoid pathway, similar to what was inferred in the original paper (Walton et al., 2016). Walton et al. reported a reduction of concentration of flavanone, p-coumaroyl hexose, quercetin glucoside, naringenin, and kaempferol glucoside for the *max2–1* mutant. Through flux-sum variability analysis, a representative of steady state metabolite pool size, the maize root model also predicted shrinkage of metabolite pool sizes of these particular metabolites. In addition, the model predicted reduced flux in some of the amino acid metabolism such as, valine-leucine-isoleucine metabolism, alanine-aspartate-glutamate metabolism, arginine metabolism, phenylalanine-tyrosine-tryptophan metabolism, and cysteine-methionine metabolism of maize root. 2,4-Dihydroxy-7-methoxy-1,4-benzoxazin-3-one-glucoside (DIMBOA-glucoside) production in benzoxazinoid pathway also showed decreased flux. DIMBOA is a tryptophan derived heteroaromatic metabolites with benzoic acid moieties that are produced in large quantities by maize roots (Cotton et al., 2019). Previous studies have showed that benzoxazinoids and their breakdown products are biocidal to some soil-borne pathogenic bacteria and fungi (Cotton et al., 2019). Hence, the lack of strigolactones in of maize root may result in weaker defence against some soil-borne pathogens. In addition, galactose metabolism, glutathione metabolism, and purine metabolism showed reduced flux. Interestingly, steroid metabolism in the *rac*-GR24 simulated maize root showed increased flux which can usually be linked to plant growth, reproduction, and responses to various abiotic and biotic stresses (Vriet et al., 2012). Fig. 2 summarizes the validation study.

**Fig. 2:**
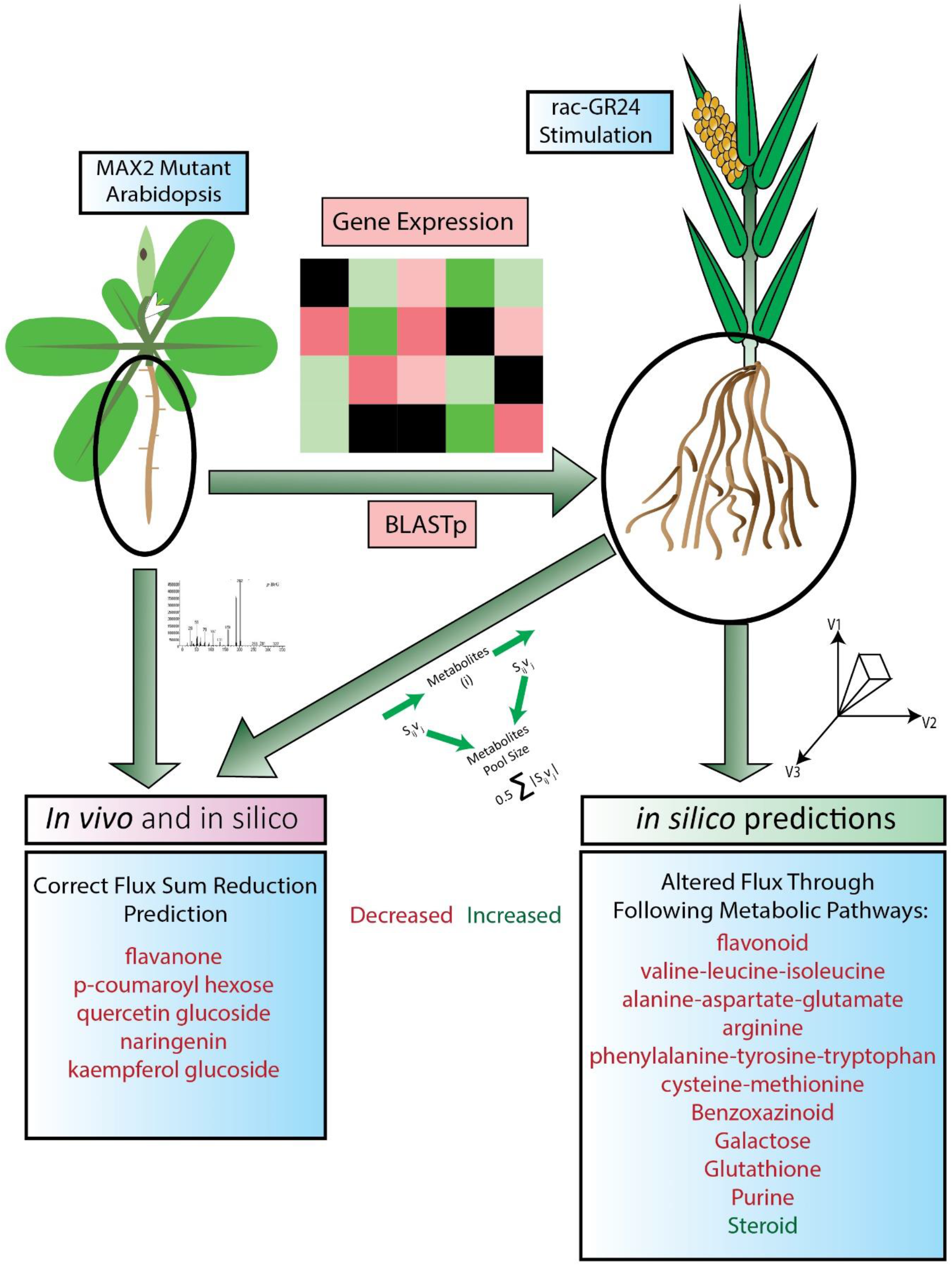
Summary of validation of the maize root GSM on the effect of strigolactone on maize root. Detail information regarding each step of validation can be found in Supplementary Information Notes S2.

### Improved model predictions through incorporation of omics data

In order to accurately model the N^−^, transcriptomics and metabolomics datasets were generated from the maize plants grown in a hydroponic system under N^−^ and N^+^. This transcriptomics data was incorporated into the root model to study metabolic reprogramming of the maize root under N^−^. Before the integration of transcriptomics data, the GSM predicted a 21% reduction of biomass in N^−^ compared to the N^+^, whereas upon the integration of transcriptomics data the trend reversed (with 285% increase in N^−^). For maize, it was reported that low N content in the growth medium increased primary root growth at the vegetative state (Mager & Ludewig, 2018). In N^−^, root tends to go deeper in the medium to scavenge more N (Cai et al., 2012). The increased growth of the root in N^−^ comes at the expense of decreased shoot growth (Mardanov et al., 1998; Puig et al., 2012). Similar root growth behavior in N^−^ was observed in other plants including *Arabidopsis thaliana* (Oldroyd & Leyser, 2020) and *Oryza sativa L. ssp. japonica* (Cai et al., 2012).

To gather further inferences on the increased root biomass production in N^−^, flux-sum variability analysis, for all the biomass metabolites was performed. Except for lipids of glycerolipid metabolism, all other metabolites showed increased but non-overlapping metabolic pool-size under N^−^ (Fig. 3). To further investigate the flux-sum variability of those lipid metabolites under N^−^, biomass production in the model was fixed between its minimum and maximum values. With the reducing biomass growth, the flux-sum ranges, of different lipid metabolites in N^−^ approached the corresponding flux-sum ranges in N^+^. When the biomass production under N^−^ was fixed to the biomass production in N^+^, most of the lipid showed similar flux-sum variability ranges as in N^+^ except for 160PC (16:0 phosphatidyl choline), 160PG (16:0 phosphatidyl glycerol), 181PC (18:1 phosphatidyl choline), 181PG (18:1 phosphatidyl glycerol), 182PC (18:2 phosphatidyl choline), 182PG (18:2 phosphatidyl glycerol), 183PC (18:3 phosphatidyl choline), and 183PG (18:3 phosphatidyl glycerol). To investigate further on these eight metabolites flux-sum variability was conducted for N^−^ with zero biomass requirement which revealed zero flux-sum for all the phosphatidyl metabolites except for those eight metabolites. Hence, these eight types of lipids were unlikely to be coupled with biomass production. Among those lipids, four of them (160PG, 181PG, 182PG, and 183PG) do not contain any nitrogen, while the rest contain nitrogen. A close observation of the glycerolipid metabolism pathway revealed that S-adenosyl-L-methionine from cysteine-methionine metabolism and citicoline from glycerophospholipid metabolism are the precursors of these four metabolites. S-Adenosyl-L-methionine is directly produced from L-methionine and the flux-sum variability comparison with the metabolomics data revealed an increased fold change of L-methionine in the N^−^. Citicoline is produced from the choline phosphate in glycerophospholipid metabolism and exhibited a widened reaction flux under N^−^. Hence, increased fold change of L-methionine and widened reaction flux for citicoline production played an important role for an elevated flux-sum ranges of 160PC, 181PC, 182PC, and 183 PC in N^−^.

**Fig. 3:**
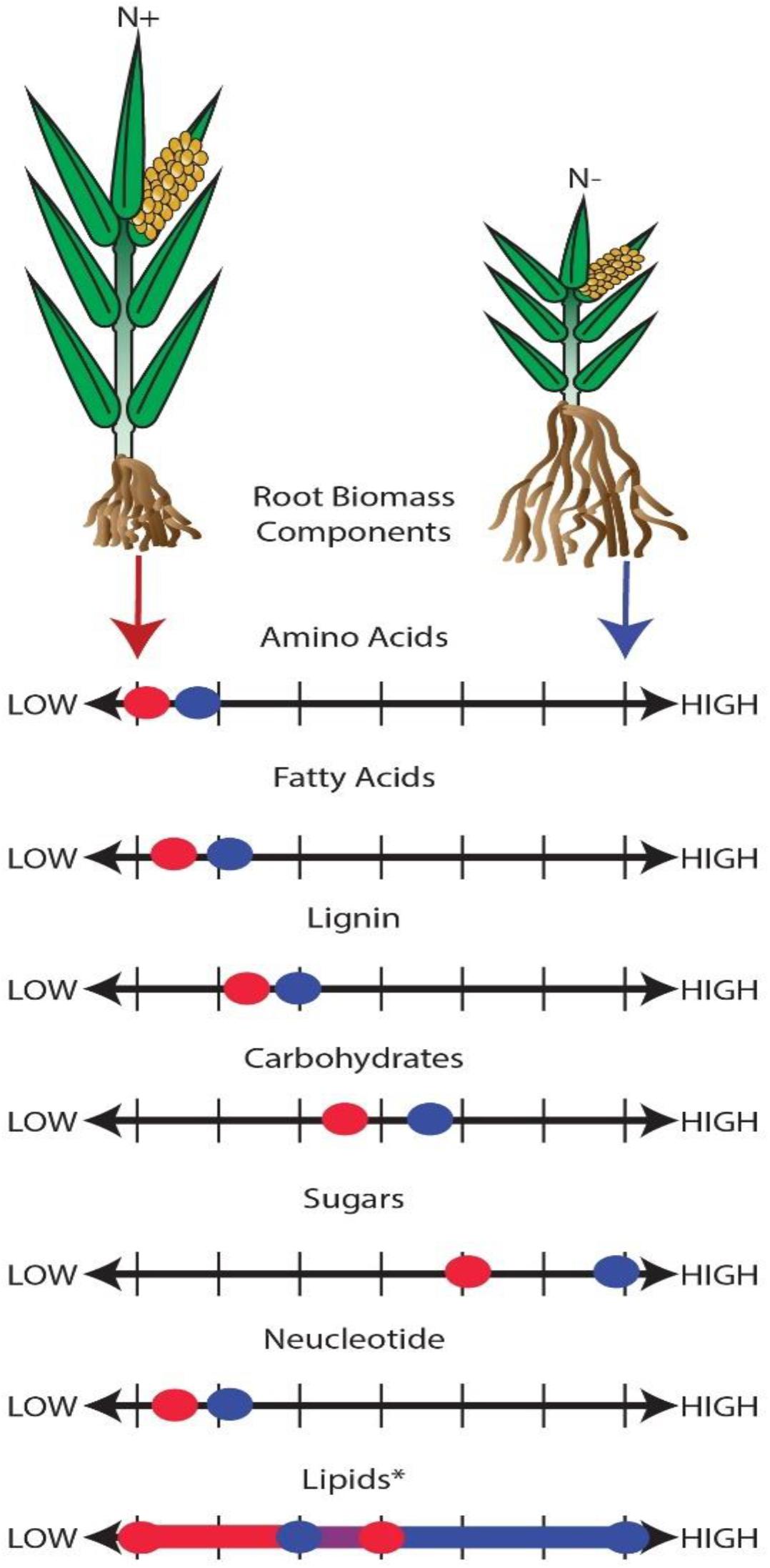
Flux-sum variability of different biomass components under N^−^ and N^+^ feeding conditions assuming maximization of biomass production. Supplementary Information Table S3 contains the full list of biomass precursors including references. Here, the size of the circle represents the flux-sum value, and the horizontal arrow represents the flux-sum range. Flux-sum variability of amino acids, fatty acids, lignin, carbohydrates, sugar, nucleotide shrunk to a point, while only for lipids, flux-sum variability showed an overlapping range. *Detail analysis of lipids metabolism is presented in Supplementary Information Notes S3.

After investigating biomass production in both conditions, the increasing or decreasing trend of metabolite content in N^−^ with respect to that measured in N^+^ was qualitatively compared with the changes in the flux-sum ranges, a representative of steady state metabolite pool size, as determined by the model. To this end, a flux-sum variability was performed, and the flux-sum ranges that did not overlap between these conditions were analyzed. An increase/decrease in the flux-sum of a metabolite between the N^−^ and the N^+^ was compared with the changes in metabolite concentration. After integration of transcriptomics data, the model was able to predict increased fold changes of metabolites between the N^−^ and N^+^. Fig. 4 demonstrates the importance of incorporating transcriptomic data as additional parameters in the model. In N^−^, the accuracy increased by more than ten-fold when the flux constraints based on transcriptomics data were incorporated to the model.

**Fig. 4:**
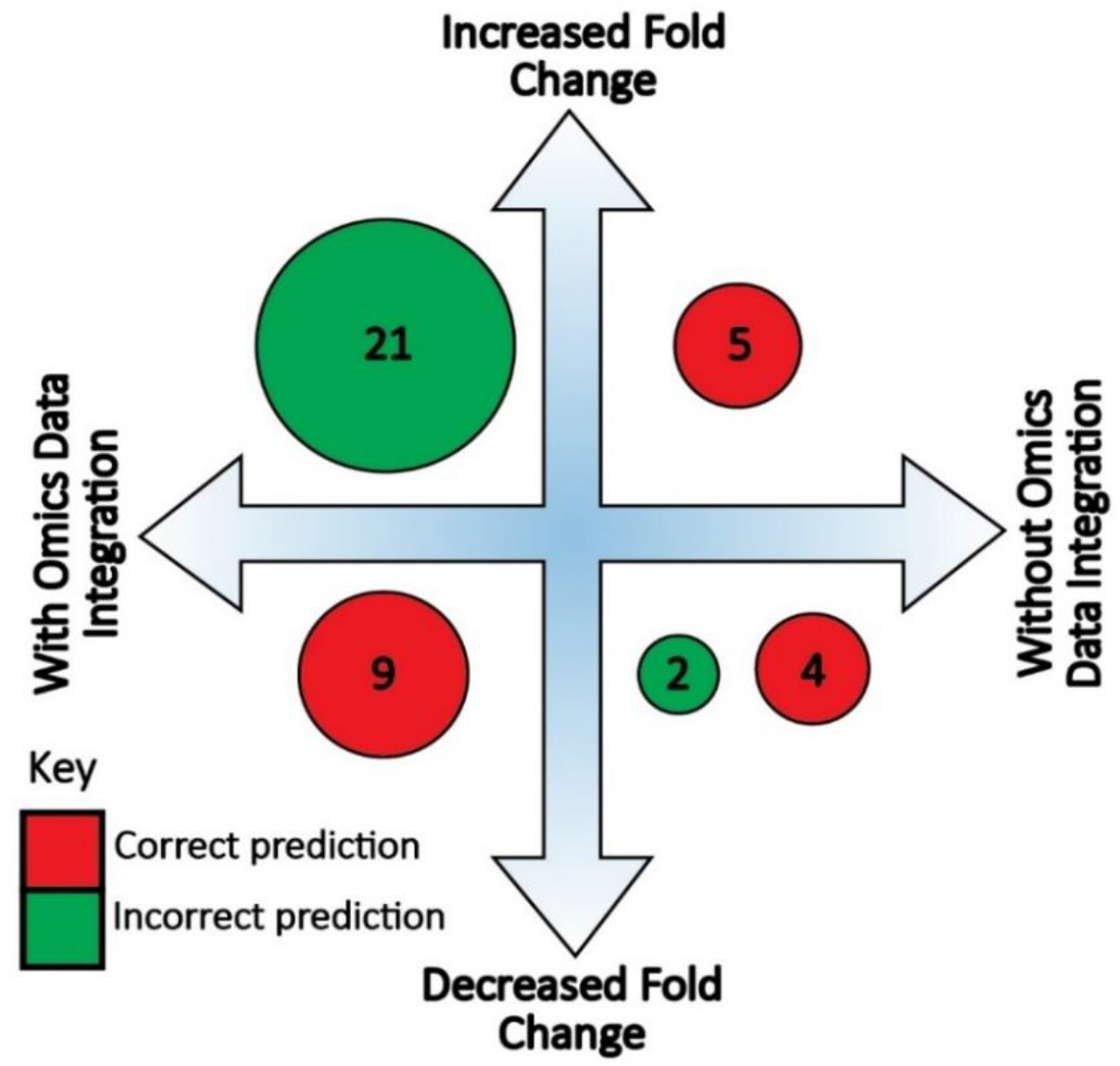
Effect of omics-based data integration on the flux-sum prediction compared with the experimental trend in metabolite concentration. The accuracy in predicting the increasing (up arrow) or decreasing (down arrow) trend in metabolite change between N^+^ and N^−^ is displayed.

Before and after the integration of transcriptomics data, five metabolites (L-lysine, glycerol-3-phosphate, glycerol, L-asparagine, and linoleic acid) were common to both feeding conditions in terms of predicting the fold change in their content. Among those metabolites, the model prediction for the L-lysine content was at the opposite of that obtained after integrating the transcriptomics-based constraints. For L-asparagine, we also found an opposite situation. For three other metabolites, the prediction remained incorrect both before and after the integration of transcriptomics data. L-asparagine is widely known as a major N carrier in plants in a number of key biological processes such as germination, vegetative growth, senescence, seed filling. Initially in the model, both the root biomass and the pool size of L-asparagine were higher in N^+^ compared to N^−^. Furthermore, after incorporation of transcriptomics data, both parameters were predicted to be higher in N^−^. Hence, the model predicted that the pool size of L-asparagine is one of the key indicators for predicting root biomass production in N^−^. Such an accumulation of L-asparagine was also observed under other abiotic stress conditions in wheat (Naidu et al., 1991, Oddy et al., 2020), Coleus (Gilbert et al., 1998), and barley (Yamaya & Matsumoto, 1989).

Before integration of transcriptomics data, the decrease of glycine in N^−^ was correctly predicted. Following the integration of transcriptomics data, the flux-sum range of glycine in N^+^ and in N^−^ overlapped with each other. In this study, we found that glycine translocation in the root through vascular tissues played an important role in this overlap. If we refer to the work Yesbergenova-Cuny et al., (2016), it has been found that glycine and arginine represented each 3% of the total phloem sap amino acid content. Thus, under N^−^, once the flux of glycine through vascular tissue was set to 85% of that of arginine, we observed that the metabolite pool size of glycine no longer overlapped between the N^+^ and the N^−^, leading to a correct prediction from the model after transcriptomics data integration. Glycine is thus an example for which transcriptomics data integration in the model can lead to erroneous flux prediction.

By incorporating transcriptomics data, the fold change in the content of several key components of the TCA cycle such as 2-oxoglutarate, succinate, fumarate, and malate were correctly predicted. 2-oxoglutarate is a key intermediate of the TCA cycle as it is the main provider of carbon skeleton to the ammonium-assimilatory pathways leading to the synthesis glutamine and glutamate (Huergo & Dixon, 2015). Other metabolites of the TCA cycle, such as succinate plays an important role during the process of symbiotic atmospheric N_2_-fixation (Flores-Tinoco et al., 2020), whereas fumarate works as a C sink for both phenylalanine and tyrosine in higher plants (Hockin et al., 2012). In root nodules, malate is the primary substrate for bacteroid respiration, thus providing energy to sustain the activity of the nitrogenase enzyme and thus the rate of N_2_-fixation. Furthermore, the fold change of several key amino acids such as lysine, arginine, methionine, cysteine, leucine, histidine, and valine that are essential for plant growth and development were correctly predicted by the model (Fig. 5). In addition, the model predicted an increase in the cholesterol and L-pipecolate contents, both being key components of root biomass production, notably under N^−^. Overall, following transcriptomics-data integration, true prediction of metabolite fold change increased from 18% to 70%. In addition, the false prediction rate of metabolite fold change decreased from 82% to 30%. Besides these correct predictions, the metabolite pool size for O-acetyl-L-serine and urea were predicted incorrectly by the model even after the integration of transcriptomics data. For these two metabolites, flux-sum range overlapped between the N^−^and the N^+^, leading to the impossibility to predict changes in their content before transcriptomic data integration. Even after transcriptomic data integration, we were not able to predict changes in their abundance. Although, we found that O-acetyl-L-serine was predominantly transported from leaves to the roots and that a minimum flux of O-acetyl-L-serine was necessary to sustain maximum biomass production in N^−^, the model prediction remained incorrect. A similar situation occurred for urea, where at maximum biomass production, when the reaction flux of arginine to urea production was reduced by 50%, the model prediction was still incorrect. Missing pathways or reactions, lack of regulatory constraints, and inadequate flow regulations through the vascular tissue under N^−^ could explain why these predictions were incorrect.

**Fig. 5:**
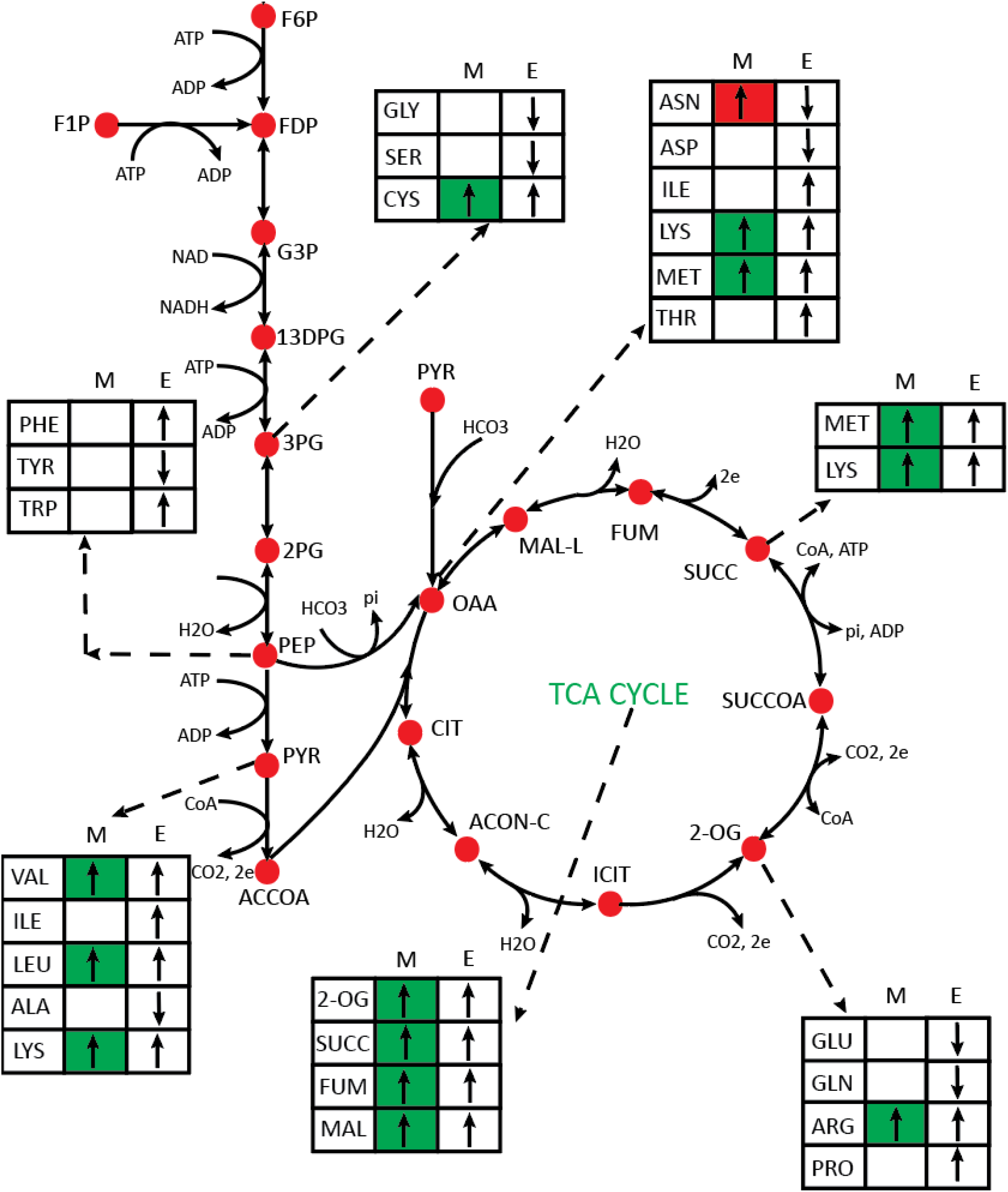
Prediction from the model for the production of carbon skeletons and the subsequent synthesis of key amino acids in central carbon metabolism. On each panel, the first column with letter indicates different amino acids or metabolites from the TCA cycle. **M** indicates the prediction from model and **E** indicates result from the experimental study. Upward arrows indicate increased metabolic pool size of a metabolite in the N^−^ compared to the N^+^. Downward arrows indicate decreased metabolic pool size of a metabolite in N^−^ compared to N^+^. The standard abbreviations for the different carbon and nitrogen metabolites were used. Flux sum level in each condition can be found in Supplementary Information Table S4.

### Flux range variations under nitrogen stress condition in hydroponic system reveals important metabolic reprogramming

Upon confirming that the model can capture the aggregate metabolic variations, we have investigated the metabolic reprogramming under N^−^. To this end, assuming that root biomass is maximized, the flux range of a specific reaction in N^−^ was compared with the flux range in N^+^ in order to study the metabolic reprogramming when plant were N-deficient.

Fig. 6 shows the flux-range of different reaction in different metabolic pathways. Central C-metabolism (CCM) plays an important role in metabolic network and is composed of the flow of C from nutrients in the different cell types via the vascular tissues to build the important components of root biomass production. The main pathways of the CCM are glycolysis/gluconeogenesis, pentose phosphate (PP) pathway, and the tricarboxylic acid (TCA) cycle. Fig. 5 shows how different metabolites from these pathways work as a precursor for several biomass components. In glycolysis, the linear pathway from glyceraldehyde-3 phosphate to acetyl-CoA showed an elevated reaction flux except for the shrinkage in the reaction flux for the interconversion between glyceraldehyde-3 phosphate and glycerone phosphate. In the PP pathway, breakdown of ribose-5 phosphate to ribose, glycerate-3 phosphate production from glyceraldehyde-3 phosphate, and production of gluconate-6 phosphate from glucose-6 phosphate exhibited widened reaction flux. In the TCA cycle, most of the reactions showed a widened reaction flux which was consistent with the experimental metabolomics data and metabolite pool-size calculations showed in Fig. 5. In order to have a perspective on the energy metabolism, flux-sum variability of ATP was performed showing that its increase in the N- was well correlated with these increased fluxes through the TCA cycle. Since CCM metabolites work as precursor to produce molecules that play a seminal regulatory role under abiotic stress conditions, in the subsequent sections we will discuss metabolic reprogramming of different pathways under N^−^.

**Fig. 6:**
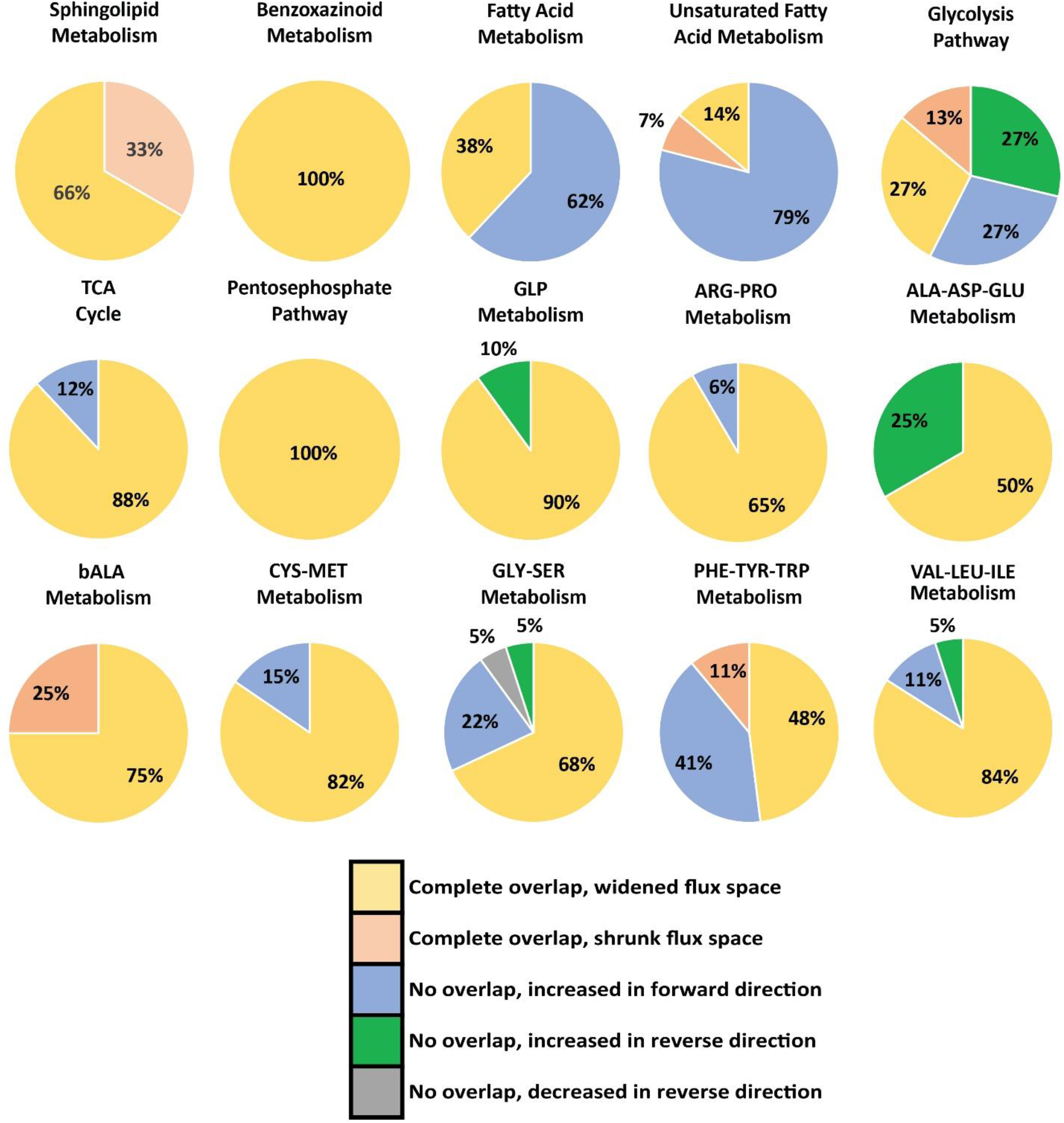
Flux range of different metabolites for the N^−^ when the N^+^ was used as a reference. The associated numbers represent the percentage of overall reactions of a specific pathway falling in each of the categories listed above.

Plant sphingolipid metabolites have important roles as a signaling molecules during both biotic and abiotic stresses (Ali et al., 2018). Sphingolipid metabolism starts by combining L-serine and palmitoyl-CoA to produce 3-dehydro-D-sphinganine. Under N^−^, the flux going through the reaction catalysed by the sphingosine kinase enzyme, which produces sphingosine 1-phosphate, increased. Sphingosine 1-phosphate plays a critical role as a signaling molecule during different stress conditions, by regulating cell growth, and suppressing programmed cell death (Spiegel & Milstien, 2003). In contrast, the interconversion between lactosylceramide to glucosylceramide exhibited a reduced flux in N^−^. Similar to sphingolipid metabolism, the reaction flux producing the final product of benzoxazinoid biosynthesis, DIMBOA, increased in N^−^. DIMBOA acts as a signaling molecule to attract the plant growth-promoting rhizobacteria Pseudomonas putida KT2440 (Costa-Gutierrez et al., 2020; Neal et al., 2012; Niemeyer, 2009). Such signaling role could also play important role during root growth in N^−^.

In fatty acid metabolism, the concentration of triacylglycerol and free fatty acids increased in Arabidopsis under N^−^ (Gaude et al., 2007). Fatty acid accumulation was also observed in other photosynthetic organisms such as *Chlamydomonas reinhardtii* (James et al., 2011; Wase et al., 2014), *Prochlorococcus marinus* (Tolonen et al., 2006), and *Auxenochlorella protothecoides* (Andeden et al., 2020) under N^−^. We also observed an accumulation of fatty acid in maize roots under N-. The flux of reactions producing octadecanoic acid, hexadecanoyl-CoA, dodecanoic acid, octanoic acid, and hexadecenoic acid significantly increased under N^−^. Similar pattern of accumulation was observed for different unsaturated fatty acids (e.g., icosenoic acid, icosadienoic acid, behenic acid, nervonic acid, arachidic acid, and lignoceric acid). To facilitate growth at the vegetative stage of plant development, plant maintain a certain stoichiometry in the C, nitrogen, and phosphorus content (Ye et al., 2014). This stoichiometry represents an optimal incorporation of macro-nutrients in order to produce biomass. However, if there is a shortage of N, C cannot be incorporated into N containing-molecules to sustain root growth and is thus stored as fatty acid when C:N cellular ratio is higher (Wase et al., 2014). Hence, accumulation of fatty acid in maize roots appear to be an important metabolic signature representative of N^−^.

Several amino acids act as precursors for the synthesis of important plant secondary metabolites and signaling molecules which play important physiological roles in a number of abiotic stresses. These signaling molecules such as polyamines are derived from arginine (Alcázar et al., 2006). Proline is another type of molecule that accumulates acting as an osmoprotectant or a compatible solute during abiotic stresses (Per et al., 2017). The family of amino acids deriving from aspartate are also involved in the energy production under abiotic stress conditions (Kirma et al., 2012). In plants, where hormone levels are modified under various abiotic stresses, cysteine acts as one of the essential precursors (Amir, 2010). Similarly, other amino acids such as lysine plays an important role as a precursor for the synthesis of a number of metabolites involved in immune signalling when there is an abiotic stress (Chen et al., 2018; Hartmann et al., 2018) and glycine, serine, and threonine are involved in phospholipid synthesis (Wattenberg, 2021). A broad spectrum of secondary metabolites with multiple biological function are further derived from the aromatic amino acids, such as phenylalanine, tyrosine, and tryptophan or from intermediates of their synthesis pathways (Tzin & Galili, 2010). Plants have evolved different strategies to minimize the adverse effects of abiotic stress conditions and several of them are connected to amino acid metabolism (Hildebrandt, 2018). A general accumulation of amino acids was observed in different plants exposed to abiotic stresses (Aleksza et al., 2017; Batista-Silva et al., 2019; Ferreira Júnior et al., 2018; Huang & Jander, 2017; Lugan et al., 2010). In this study, an accumulation of amino acid was also observed under N^−^. Overall, under N^−^, flux of all reactions involved in arginine-proline, alanine-aspartate-glutamate, cysteine-methionine, valine-leucine-isoleucine, and histidine biosynthesis showed elevated reaction flux. In the β-alanine pathway, under N-, most of the reactions showed elevated reaction flux except for the degradation of 3-oxopropanoate to acetyl-CoA, exhibited reduced reaction flux. As acetyl-CoA is the precursor of fatty acid biosynthesis, to meet the increasing fatty acid demand, an increased dissociation of malonyl-CoA to acetyl-CoA during β-alanine biosynthesis was observed. In the glycine-serine-threonine biosynthesis pathway, similar to β-alanine, most of the reactions showed elevated fluxes under N^−^. Interestingly, under N^−^, interconversion between glycine and threonine was in favour of threonine production. The experimental metabolomics data also showed that there was an increase in threonine compared to that observed for glycine (Fig. 5). Similarly, in the phenylalanine-tyrosine-tryptophan biosynthesis pathway, most of the reactions showed elevated flux except for the reactions in the linear pathway, producing chorismite from shikimic acid.

In plants, increased starch accumulation in the leaves is another metabolic reprogramming observed under N^−^, such as duckweed (Yu et al., 2017), Arabidopsis (Krapp et al., 2011), and maize (Amiour et al., 2012). In this study, a similar pattern of starch accumulation in the roots was observed. Reaction flux from amylose to starch showed elevated flux. Similarly, the dissociation of starch to α-D-glucose-1 phosphate showed reduced flux under N^−^. Starch functions as one of the largest sources of C sink in a plant. During vegetative growth, the roots and immature leaves are the largest carbon sinks. Then, following the transition to reproductive growth, floral organs, reproductive and storage organs become the largest sinks for carbon. However, if nutrients become limited at any stage of the life cycle, more C is allocated to the roots in order to increase soil mineral acquisition, resulting in a shift in the relative sink balance and C partitioning within the plant (Eghball & Maranville, 1993). In N^−^, a similar pattern of partitioning was observed for other C sinks such as lignins, where the linear pathway from p-coumaric acid to the p-hydroxy-phenyl lignin showed an increased flux. Overall, these metabolic reprogramming under N^−^ provided important insights relating the phenotypic changes of the roots to the underlying metabolism.

In this work, a GSM for maize root was reconstructed which, upon incorporation of omics data, revealed important metabolic reprogramming under N^−^. The reconstruction of maize root GSM predicted the increased root biomass production under N^−^. Beside predicting important metabolic reprogramming in CCM, FAM, amino acid metabolism, and several other secondary metabolisms, maize root GSM also revealed that several metabolites, such as L-methionine, L-asparagine, L-lysine, cholesterol, and L-pipecolate, were important compounds involved in root biomass production. Furthermore, this study revealed eight phosphatidyl choline and phosphatidyl glycerol metabolites, not directly coupled with biomass production, played an important role in root growth under N^−^.

Future research will be focused on the reconstruction of tissue-specific models for kernel, stalk, and tassel. Then, this root GSM will be combined to those tissue-specific models and the previously reconstructed maize-leaf model (Simons et al., 2014) to develop a whole plant maize GSM. A whole plant GSM will be useful to elucidate the flow of different micro and macro nutrients from the root to the shoots and to the reproductive organs in a maize plant. Furthermore, whole plant maize GSM, will allow to study metabolic reprogramming under various other stress conditions such as phosphate deficiency, salinity, draught and thermal stresses, heavy metal accumulation, elevated levels of CO_2_, and in the presence of beneficial soil microorganisms. Such studies will allow to identify key metabolic pathways and markers representative of these stresses, which can potentially be used to select maize genotypes adaptive to diverse favorable or unfavorable environmental conditions.

## Acknowledgments

The authors thank Mohammad Mazharul Islam for constructive comments on the manuscript. We also thank Drs. Gilles Clément, Gregory Mouille, and Martine Miquel for performing the metabolome and some of the biomass component analyses which were carried out at the Obervatoire du Végétal Chimie Metabolisme of the Institute Jean-Pierre Bourgin, INRA, Versailles-Grignon. We gratefully acknowledge funding support from National Science Foundation (NSF) CAREER grant (25-1106-0039-001) and The Center for Bioenergy Innovation, a U.S. Department of Energy Research Center supported by the Office of Biological and Environmental Research in the DOE Office of Science.

## Author Contribution

RS, CDM, and BH conceived the study; RS supervised the study; NA, IQ, and BH performed and analyzed the experimental studies; NBC performed all in silico experiments and analyses; NBC, WS, and DS developed the required software programs, models, and graphics; NBC. wrote the original manuscript and RS, BH, and CDM. edited the manuscript. All authors have reviewed and approved the submission of the manuscript.

## Data availability

The data that support for the findings of this study can be found in the related cited articles and/or in the supplementary materials.

## Supporting Information

**Notes S1** Detail experimental procedure

**Table S1** Genome-Scale Metabolic Model of Maize Root

**Table S2** Experimental transcriptomics and metabolomics data

**Notes S2** Arabidopsis validation

**Table S3** Root biomass information

**Notes S3** Lipid analysis

**Table S4** Flux-sum level in each condition

